# Python metabolomics uncovers a conserved postprandial metabolite and gut-brain feeding pathway

**DOI:** 10.64898/2026.01.09.698526

**Authors:** Shuke Xiao, Mengjie Wang, Thomas G. Martin, Barry Scott, Xing Fang, Xinming Liu, Yongjie Yang, Sipei Fu, Steven D. Truong, Jack F. Gugel, Gregory L. Maas, Marcus P. Mullen, Jennifer Hampton Hill, Veronica L. Li, Andrew L. Markhard, Mingming Zhao, Wei Qi, Saranya C. Reghupaty, Meng Zhao, Jan Spaas, Wei Wei, Trine Moholdt, John A. Hawley, Christian T. Voldstedlund, Erik A. Richter, Xiaoke Chen, Katrin J. Svensson, Daniel Bernstein, Leslie A. Leinwand, Yong Xu, Jonathan Z. Long

## Abstract

Most mammals consume small and frequent meals. By contrast, pythons are ambush predators that exhibit extreme feeding and fasting patterns and provide a unique model for uncovering molecular mediators of the postprandial response^1–3^. Using untargeted metabolomics, here we show that circulating levels of the metabolite *para*-tyramine-O-sulfate (pTOS) are increased >1,000-fold in pythons after a single meal. In pythons, pTOS production occurs in a microbiome-dependent manner via sequential decarboxylation and sulfation of dietary tyrosine. In both pythons and mice, pTOS administration activates a neural population in the ventromedial hypothalamus (VMH). In mice, these VMH neurons are required for the anorexigenic effects of pTOS. Chronic administration of pTOS to diet-induced obese male mice suppresses food intake and body weight. pTOS is also present in human blood, where its levels are increased after a meal. Together, these data uncover a conserved postprandial anorexigenic metabolite that links nutrient intake to energy balance.

## Main Text

Organismal energy homeostasis is a highly dynamic process that is strongly influenced by nutrient availability. Nutrient consumption induces a postprandial phase marked by the release of neuroendocrine signals and hormones, such as insulin, that promote anabolism and nutrient storage. Conversely, in the fasted state, the release of other metabolic messengers, such as glucagon, induce a shift towards mobilization of internal energy reserves. To date, much of our understanding about the endocrine control of energy homeostasis during feeding or fasting comes from studies in mammals such as humans and rodents. While informative, mammals are nevertheless adapted to consuming small (<1-2% of body weight) and frequent (1-3 times per day) meals. This physiology places a fundamental limit on the degree to which fluctuations of energy states and postprandial physiologic responses can occur.

By contrast, the Burmese python (*Python molurus bivittatus*) offers a striking example of an organism that demonstrates extreme feeding and fasting patterns, as well as postprandial physiology^1–3^. Burmese pythons are ambush predators that can fast for extended periods, sometimes up to 12 to 18 months. When they do feed, they can consume prey equal to their own body weight in a single meal^1^. Following such a meal, Burmese pythons display a remarkable array of postprandial responses, including >40-fold increase in energy expenditure, sustained tissue protein synthesis, and >50% increase in the size of most organs^4–11^. Upon completion of digestion, many of these responses revert to levels close to that of the original fasted state^1,2,11–13^, demonstrating that these snakes undergo an extreme and reversible postprandial responses with every meal. Beyond Burmese pythons, many other members of the Pythonidae family also exhibit sit-and-wait predatory lifestyles and extreme feeding and fasting patterns, including ball pythons (*Python regius*), African rock pythons (*Python sebae*), and reticulated pythons (*Malayopython reticulatus*)^11,14^.

We hypothesized that the extreme feeding patterns and physiologic responses of pythons would also be mirrored by equally extreme molecular responses. The python could therefore provide a unique opportunity to discover postprandial molecules that would otherwise be missed in more classical organisms, such as mammals, which have more limited fluctuations in their energy states and postprandial physiology. Such molecules would not only be most readily detectable in pythons would likely be potentially informative for mammalian physiology as well.

To date, only a limited number of previous efforts have sought to understand the postprandially regulated molecules in such a physiologically extreme organism^4,15–17^. For instance, metabolomic studies have revealed an increase in fatty acids and bile acids in postprandial python plasma^4^, a mixture of which was shown to be sufficient to stimulate cardiac hypertrophy when administered to either pythons or mice^4^. However, these past studies have been relatively narrowly focused on a small set of predefined molecules. We reasoned that an unbiased metabolomic analysis of plasma from fasted and fed pythons might reveal heretofore unrecognized signaling molecules that play a critical role in the python postprandial response. Using untargeted metabolomics, here we identify a metabolite, *para*-tyramine-O-sulfate (pTOS), as the most dramatically induced metabolite (>1,000-fold) in postprandial python plasma. Follow up studies in both pythons and mammals show that pTOS functions as a conserved anorexigenic postprandial metabolite gut-brain signal.

## Results

### Metabolomics of postprandial python plasma

We used both targeted and untargeted metabolomics to identify circulating metabolites significantly altered by feeding in Burmese pythons. Pythons (approximately 2 years old and weighing 1.5-2.5 kg at the start of the study) were fed once every 28 days, with each meal amounting to approximately 25% of their body weight. Plasma samples were collected from fasted pythons and from pythons sacrificed three days post-feeding (3DPF, Figure 1A).

**Figure. 1.**
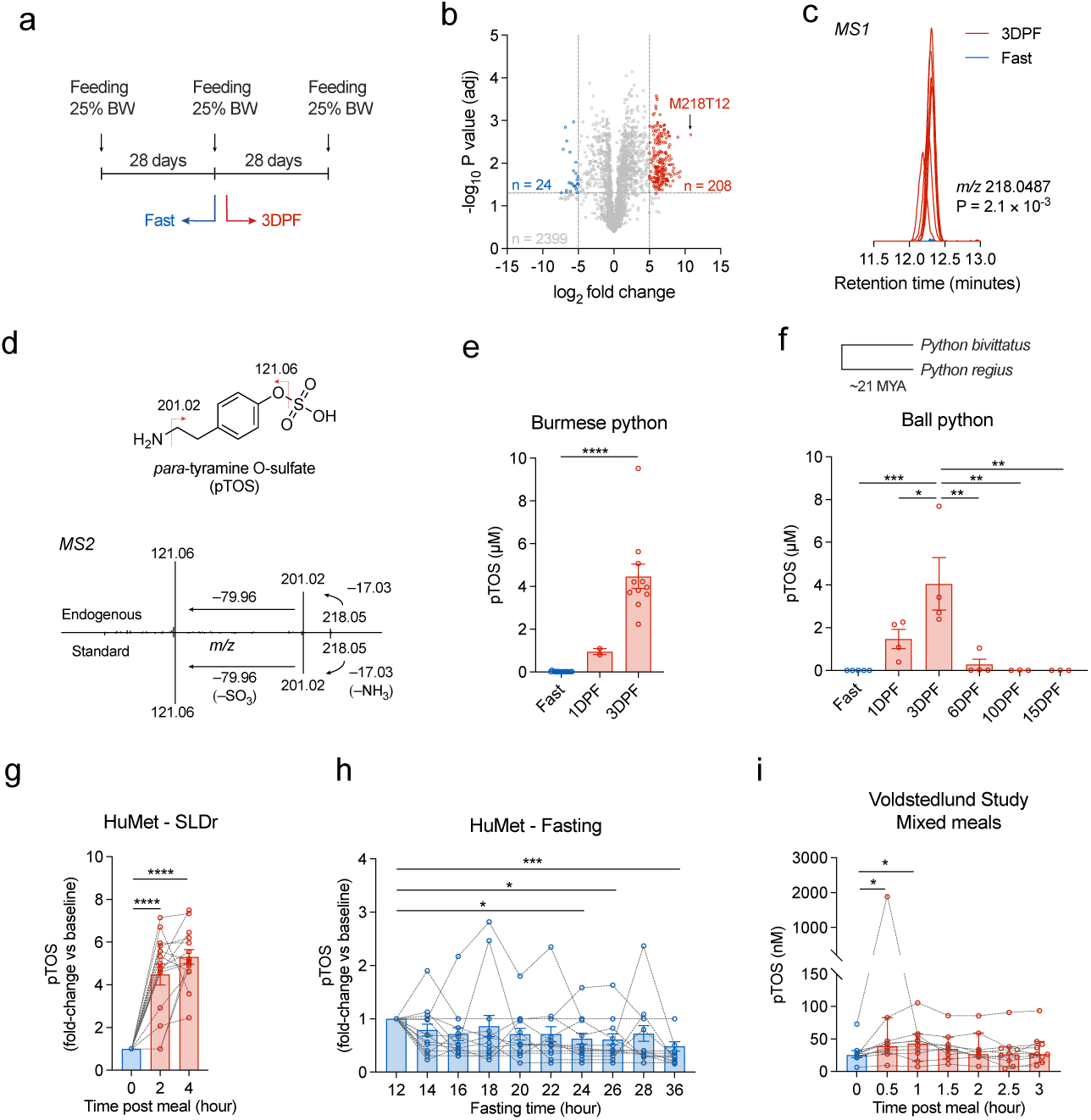
Postprandial pTOS increase in python and human plasma. (a) Feeding scheme of Burmese pythons (*Python molurus bivittatus*) and the time points of blood plasma collection. (b) Global changes in metabolites between 3DPF and 28-day fasted Burmese python plasma using untargeted metabolomics. (n = 6 per group). (c) Extracted ion chromatograms of the m/z = 218.0487 from fasting and 3DPF plasma. (d) Chemical structure of pTOS (above) and tandem mass spectrometry fragmentation of the endogenous m/z = 218.0487 and authentic pTOS standard (below). (e, f) Absolute quantitation of pTOS in Burmese pythons (e) and in ball pythons (*Python regius*) at the indicated time point (f). (g) Relative changes of pTOS levels in human plasma with the recovery standardized liquid diet (SLDr) meal in the “HuMet” study (n = 15 human subjects). (h) Relative changes of pTOS levels in human plasma with fasting in the “HuMet” study (n = 15 human subjects). (i) Absolute quantitation of pTOS in human plasma after three mixed meal tests in the “Voldstedlund” study (n = 10 human subjects). In (e), n = 15 for Fast group, n = 2 for 1DPF group, n = 11 for 3DPF group. In (f), n = 5 for Fast group, n = 4 for 1DPF, 3DPF, 6DPF groups, n = 3 for 10 DPF and 15 DPF groups. Data are shown as mean ± SEM in (e-h). P values in (b) and (c) were calculated with two-tailed t tests and adjusted with Benjamini-Hochberg method. P values in (e) and (f) were calculated with one-way ANOVA followed by Turkey’s multiple comparison tests. P values in (g, h) were calculated with Repeated Measures One-Way ANOVA followed by Dunnett’s multiple comparisons test. Data in (i) are shown as median ± 95% confidence intervals. P values were calculated with two-sided Friedman test followed by Dunn’s multiple comparisons test. * p < 0.05, ** p < 0.01, *** p < 0.001, **** p < 0.0001. Exact P values are as follows: (e) Fast vs 3DPF, P = 1.5 × 10^-5^; (f) Fast vs 3DPF, P = 0.0006, 1DPF vs 3DPF, P = 0.0440, 3DPF vs 6DPF, P = 0.0021, 3DPF vs 10 DPF, P = 0.0022, 3DPF vs 15DPF, P = 0.0022; (g) 0 vs 2, P = 2.5 × 10^-8^, 0 vs 4, P = 3.0 × 10^-10^; (h) 12 vs 24, P = 0.0304, 12 VS 26, P = 0.0226, 12 vs 36, P = 0.0008; (i) 0 vs 0.5, P = 0.0312, 0 vs 1, P = 0.0161.

In the comparison between 3DPF and fasted python plasma samples, our targeted metabolomics platform detected several metabolites that were increased by 4-20-fold postprandially. For instance, we observed an increase in many circulating amino acids, especially the non-essential amino acids glycine (20-fold) and proline (9-fold) (Extended Data Figure 1a). Fumarate and malate were also increased by 4-8-fold (Extended Data Figure 1b). Lastly, several species of very long-chain fatty acids, such as lignoceric acid (C24:0), hexacosanoic acid (C26:0), and hexacosenoic acid (C26:1) were also increased by 4-6-fold (Extended Data figure 1c).

In our untargeted metabolomics platform, many more metabolites with even more dramatic increases after feeding were detected. Using a cutoff of a 2^5^-fold change, we found 208 metabolite features significantly (P_adj_ < 0.05) increased, and 24 metabolites significantly decreased, in plasma from 3DPF versus fasted samples (Figure 1b, Supplementary Table 1). The metabolite feature with the greatest fold increase after feeding (>1,000-fold) in python plasma was an unknown molecule with a mass-to-charge ratio (*m/z*) of 218.0487 in positive ionization mode (P_adj_ = 2.1 × 10^-3^; Figure 1c). This exact mass enabled us to deduce a chemical formula for the parent ion of C_8_H_11_NO_4_S. Tandem mass spectrometry (MS/MS) of this feature revealed two distinct fragment ions: one at *m/z* 201.0187, indicating the loss of an NH_3_ group, and another at *m/z* 121.0634, corresponding to the neutral loss of a sulfate group (SO_3_). This C_8_H_11_NO_4_S metabolite feature was also detectable in negative ionization mode with an *m/z* of 216.0331 (Extended Data Figure 2a) and a loss of sulfate (−80, Extended Data figure 2b) upon MS/MS fragmentation. Based on the parent chemical formula and fragmentation patterns, we surmised that this metabolite contained a primary amine, a phenolic sulfate group, and two additional methylene groups. We therefore tentatively assigned the structure as *para*-tyramine-O-sulfate (pTOS; Figure 1d). This structural assignment was confirmed when the endogenous plasma peak was compared to an authentic pTOS standard generated via chemical synthesis: the authentic standard matched both the fragmentation pattern (Figure 1d; Extended Data figure 2b) and the retention time (Extended Data figure 2c) of the endogenous metabolite in both positive and negative ionization modes.

Quantitative analyses revealed that plasma pTOS concentrations increased from 20 ± 5 nM in the fasted state to 1.0 ± 0.1 µM at 1DPF and peaked at 4.5 ± 0.6 µM at 3DPF in Burmese pythons (Figure 1e). To determine the generality of postprandial pTOS increases, we also measured plasma pTOS levels in ball pythons (*Python regius*), a species that diverged from Burmese pythons approximately 21 million years ago (Figure 1f). In ball pythons, a similar postprandial rise in pTOS was observed: pTOS concentrations in the fasted state were 1.2 ± 0.6 nM, increased to 1.5 ± 0.4 µM by 1DPF, and peaked at 4.1 ± 1.2 µM at 3DPF. In ball pythons, we were able to collect a longer postprandial plasma time course and observed that pTOS levels drastically decreased to sub-micromolar levels at 6DPF, and back to nanomolar range at 10DPF and 15DPF (Figure 1f). The time course is similar (peak at days 1-3) to that previously reported for metabolic rate changes after a meal in pythons^18^.

### pTOS levels and postprandial regulation in humans and mice

pTOS is a poorly studied metabolite. In humans, sporadic reports have detected pTOS in urine as a rapidly excreted molecule^19,20^. Its presence in circulation and postprandial regulation in humans had not been rigorously studied.

We first examined public metabolomic datasets where circulating metabolites were measured before and after a meal test in humans. We identified three study cohorts and a total of six meal tests where blood pTOS measurements were available. The first “HuMet” study cohort consisted of 15 healthy male participants who were exposed to several physiologic challenges, including three standardized liquid diet (SLD) meals^21^. The first meal consisted of a recovery meal after a 36 h fast (“SLDr”); under these conditions, pTOS levels increased by 2-8-fold (Figure 1g). The other two meals also led to increases of pTOS by an average of ∼1.5-fold and ∼3-fold (Extended Data figures 3a, b). Interestingly, in the same study, pTOS levels also decreased over the initial 36 h fasting period (Figure 1h). The second study cohort (“Moholdt”) consisted of N = 24 healthy young men who followed either a habitual diet or brief high-fat diet feeding^22^. In this study, circulating pTOS levels were measured either before breakfast or after dinner and exhibited an average of ∼5-fold increase after dinner regardless of the diet (Extended Data figure 3c,d). One individual exhibited >30-fold increase in pTOS levels. The third study cohort (“Agueusop”) examined the metabolic changes pre- and one hour post- a standardized mixed meal in N = 30 individuals that were either healthy, prediabetic, or with type 2 diabetes^23^. Here, pTOS levels were not changed after the standardized mixed meal (Extended Data figure 3e). Therefore, across the six available human meal-test datasets, five showed that pTOS increases after a meal. The single exception involved participants with prediabetes or type 2 diabetes. We conclude that pTOS is present in the circulation in humans. In addition, a modest postprandial induction of pTOS in humans is generally observed across most studies but may be absent in participants with impaired glucose homeostasis.

To independently verify postprandial induction of pTOS in humans, and to quantitatively determine absolute concentrations of pTOS in circulation, we used LC-MS to measure pTOS levels from blood samples that had been previously collected during a meal test led by Voldstedlund et al.^24^ In this study, 10 healthy male participants were fasted for 6.5 hours and then consumed three mixed-nutrient meals (one solid, two liquid). Plasma was collected in the fasted state and at various time points after initiation of the first meal. We quantified plasma pTOS levels to be ∼25 nM in the fasted state and increased by an average of ∼2-fold post-meal (Figure 1i), with one individual exhibiting a >25-fold increase of pTOS and circulating concentrations of ∼2 µM at 0.5 h post-meal.

Lastly, we examined circulating pTOS levels in mice. Surprisingly, we did not robustly detect pTOS in mouse plasma either in the basal state or after a meal (Extended Data figure 3f, limit of detection < 1 nM). We also surveyed publicly available datasets of human, mouse, or rat blood metabolomics. pTOS was reported in 20 of 537 human studies (4%) but in 0 of 237 mouse and 0 of 52 rat studies (Extended Data figure 3g). By contrast, the amino acid tyrosine was detected with equal frequency in metabolomics datasets from all three species (39% human, 34% mouse, 33% rat) (Extended Data figure 3h). We conclude that pTOS is not present, or at very low < 1 nM levels, in mouse blood.

### Python biosynthesis of pTOS from dietary tyrosine

To understand the mechanisms underlying the dramatic postprandial induction of pTOS in pythons, we sought to understand the pathways of pTOS biosynthesis. The structural similarity between pTOS and the amino acid tyrosine suggests a sequential biosynthesis pathway involving decarboxylation of the dietary amino acid tyrosine to produce tyramine, followed by sulfation to generate pTOS (Figure 2a). When we administered tyrosine to fasted ball pythons (1 g/kg, per os [p.o.]), we observed a 5.1-fold increase in plasma pTOS levels after 24 h compared to vehicle controls (Figure 2b). We conclude that dietary tyrosine can be metabolically converted to pTOS in pythons. This rise, though significant, was less pronounced than the postprandial response, likely because the extensive gastrointestinal remodeling that accompanies feeding (including increased blood flow to the gut and upregulation of amino acid transporters) likely does not occur during acute tyrosine gavage.

**Figure 2.**
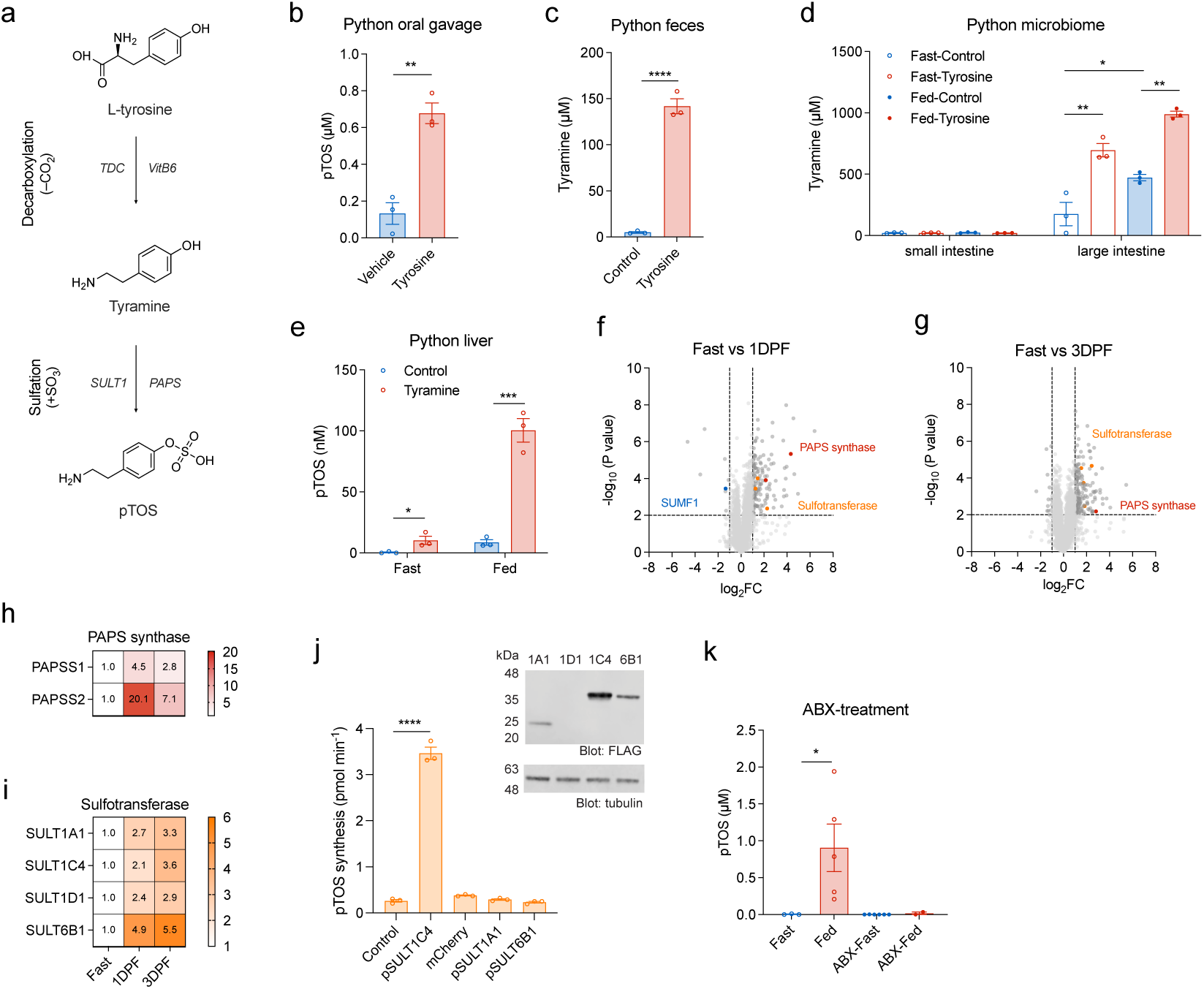
pTOS production from dietary tyrosine. (a) Scheme of the biochemical reactions involved in pTOS synthesis. Dietary tyrosine is converted to pTOS through decarboxylation and subsequent sulfation. (b) Plasma pTOS levels in fasted (28DPF) ball pythons one day after tyrosine (1g/kg) or vehicle oral gavage. (n = 3/group) (c) Tyramine levels in the media of anaerobic bacterial cultures of ball python feces using the SAAC media (control, containing 1 mM tyrosine and 0.1 mg/L vitamin B6) or supplemented with 50 mM tyrosine and 5 mg/L vitamin B6 (tyrosine). (n = 3/group) (d) Tyramine levels in the GIFU media of anaerobic cultures of the microbiome collected from the small and large intestines of fasted (107DPF) or fed (3DPF) ball pythons. (n = 3/group). (e) pTOS levels in the conditioned media of python liver slices incubated with tyramine (1mM) or vehicle for 16 hrs. Livers were freshly isolated from fasted (31DPF) or fed (3DPF) ball pythons. (n = 3/group). (f, g) Proteomics changes in livers of fasted ball pythons at the indicated time point. (h, i) Fold changes of PAPS synthases (h) and SULTs (i) in livers at the indicated time point. Fasted liver samples were normalized to 1. In (f-i), n = 5 for Fast and 1DPF groups, n = 4 for 3DPF group. (j) Sulfation activity of HEK293T cell lysates expressing FLAG-tagged python SULTs, using tyramine as the substrate. Western blot analysis shows expression of the python SULTs (anti-FLAG, top) and loading control (anti-tubulin, bottom). (k) Plasma pTOS levels in fasted (28DPF) or fed (3DPF) Burmese pythons following antibiotics (ABX) treatment. Vehicle-treated pythons: n = 3 for Fast, n = 5 for Fed; ABX-treated pythons: n = 6 for Fast, n = 2 for Fed. TDC, tyrosine decarboxylase. SULT1, sulfotransferase 1. PAPS, 3’-Phosphoadenosine-5’-phosphosulfate. SUMF1, sulfatase-modifying factor 1. Data are shown as mean ± SEM. P values were calculated with two-tailed t tests (b, c), one way ANOVA followed by Holm-Šidák’s multiple comparisons test (d), multiple t tests (two-tailed) (e), two-tailed t tests adjusted with Benjamini-Hochberg method (f, g), one way ANOVA followed by Dunnett’s multiple comparisons test (j), one way ANOVA followed by Bonferroni’s multiple comparisons test (k). * p < 0.05. **p < 0.01, **** p < 0.0001. Exact P values are as follows: (b) P = 0.002; (c) 7.5 × 10^-5^; (d) Large intestine, Fast-Control vs Fast-Tyrosine, P = 0.001, Fast-Control vs Fed-Control, P = 0.0192, Fed-Control vs Fed-Tyrosine, P = 0.001; (e) Fast, P = 0.0424, Fed, P = 0.0007; (j) Control vs pSULT1C4, P = 2.1 × 10^-11^; (k) Fast vs Fed, P = 0.0234.

We next focused on each step of the tyrosine-to-pTOS pathway. The first tyrosine decarboxylation step can be catalyzed by several enzymes, the most well-studied of which are bacterial tyrosine decarboxylases (TDCs; EC 4.1.1.25)^25^. These enzymes exhibit high affinity and specificity for tyrosine and are highly expressed in several gut bacterial taxa such as *Lactobacillus* and *Enterococcus* species. We therefore isolated and cultured fecal bacteria from ball pythons in standard amino acid complete (SAAC) media and examined the production of pTOS in vitro. Addition of tyrosine (50 mM) and the cofactor pyridoxal 5′-phosphate (5 mg/L) for 2 days significantly increased tyramine production in both cells and the media compared to cells that were not treated with tyrosine starting material (Figure 2c). When we further isolated and cultured bacteria from ball python small and large intestines under fasted or fed conditions, we observed exclusive tyramine production from large intestine microbes, which was further enhanced in the fed versus fasted condition (Figure 2d).

Next, we examined the second step of tyramine sulfation. Sulfotransferases (SULTs) are a class of enzymes that catalyze metabolite sulfation and exhibit liver-enriched expression^26^. We reasoned that python hepatic sulfotransferase activity may mediate conversion of tyramine to pTOS. Consistent with this premise, robust production of pTOS was observed following the addition of tyramine to liver tissue slices that had been freshly isolated from fasted pythons (Figure 2e). When similar experiments were performed using liver tissue slices isolated from 3DPF pythons, pTOS production was further increased (Figure 2e). We performed tandem mass tag (TMT)-based quantitative proteomics analysis on fasted, 1DPF, and 3DPF python liver (Figure 2f,g and Supplementary Table 2). We observed significant increases in the abundance of python hepatic enzymes involved in metabolite sulfation reactions. For instance, two 3’-phosphoadenosine 5’-phosphosulfate (PAPS) synthases, which provide the sulfur donor in these reactions, were increased by 2-20-fold in the postprandial state (Figure 2h). In addition, four sulfotransferases (SULTs) were also increased by 2-6-fold (Figure 2i). To determine if any of these python sulfotransferases could catalyze tyramine sulfation, we expressed individual python SULT enzymes in HEK293T cells and tested cell lysates for sulfotransferase activity. We confirmed expression of python SULT1A1, SULT1C4, and SULT6B1 by anti-flag Western blotting, while python SULT1D1 failed to express under these conditions. Cell lysates expressing python SULT1C4, but not SULT1A1 or SULT6B1, exhibited robust tyramine sulfation activity (Figure 2j). We conclude that a two-step biochemical pathway in pythons involving decarboxylation and hepatic sulfation converts dietary tyrosine to pTOS.

Lastly, we sought to critically test the requirement of gut bacteria in postprandial pTOS production in pythons. We therefore treated pythons with a cocktail of antibiotics (ABX) (200 mg/kg metronidazole, 200 mg/kg ampicillin, 100 mg/kg neomycin, and 100 mg/kg erythromycin) and measured pTOS levels after a meal. Antibiotic-mediated depletion of the gut microbiome was confirmed by reduced gut bacterial content (Extended Data figure 4a,b). Control pythons exhibited expected postprandial increase of pTOS (Figure 2k). This rise in pTOS was largely abolished in ABX-treated pythons (Figure 2k). We conclude that gut bacteria are required for postprandial pTOS induction in pythons.

### pTOS suppresses feeding behaviors in mice

No biological function has been attributed to pTOS. Its dramatic induction in the python postprandial phase prompted us to investigate whether pTOS might influence whole-body energy metabolism. We performed gain-of-function experiments by administering pTOS (50 mg/kg, i.p.) to male mice in metabolic chambers. Such a dose was chosen to achieve circulating pTOS levels that are comparable to postprandial python levels. Under these conditions, we observed an increase in plasma pTOS levels which peaked at 11.4 ± 1.2 µM at 1 h, decreased to 2.9 ± 0.6 µM at 2 h, and 0.5 ± 0.1 µM at 4 h (Extended Data figure 5a). Mice treated with pTOS exhibited a significant reduction in food intake (mean ± SEM, vehicle: 4.39 ± 0.28 g, pTOS: 3.58 ± 0.16 g, p = 0.024) over the 24 h period (Figure 3a,b). While the respiratory exchange ratio (RER) trended towards lower in pTOS-treated mice, this difference was not statistically significant (Figure 3c,d). It may also be possible that other, more subtle aspects of glucose metabolism are concurrently altered in pTOS treated mice, leading to increased RER above what would be expected from the reduced feeding. Energy expenditure (Figure 3e,f and Extended Data figure 5b-d) and locomotor activity (Figure 3g,h) were also not statistically different between vehicle and pTOS-treated mice. Importantly, pTOS administration did not affect water intake (Figure 3I), induce conditional flavor avoidance (Figure 3j and Extended Data figure 5e), or affect sucrose preference (Figure 3k). pTOS administration also did not alter blood glucose levels (Extended Data figure 5f).

**Figure 3.**
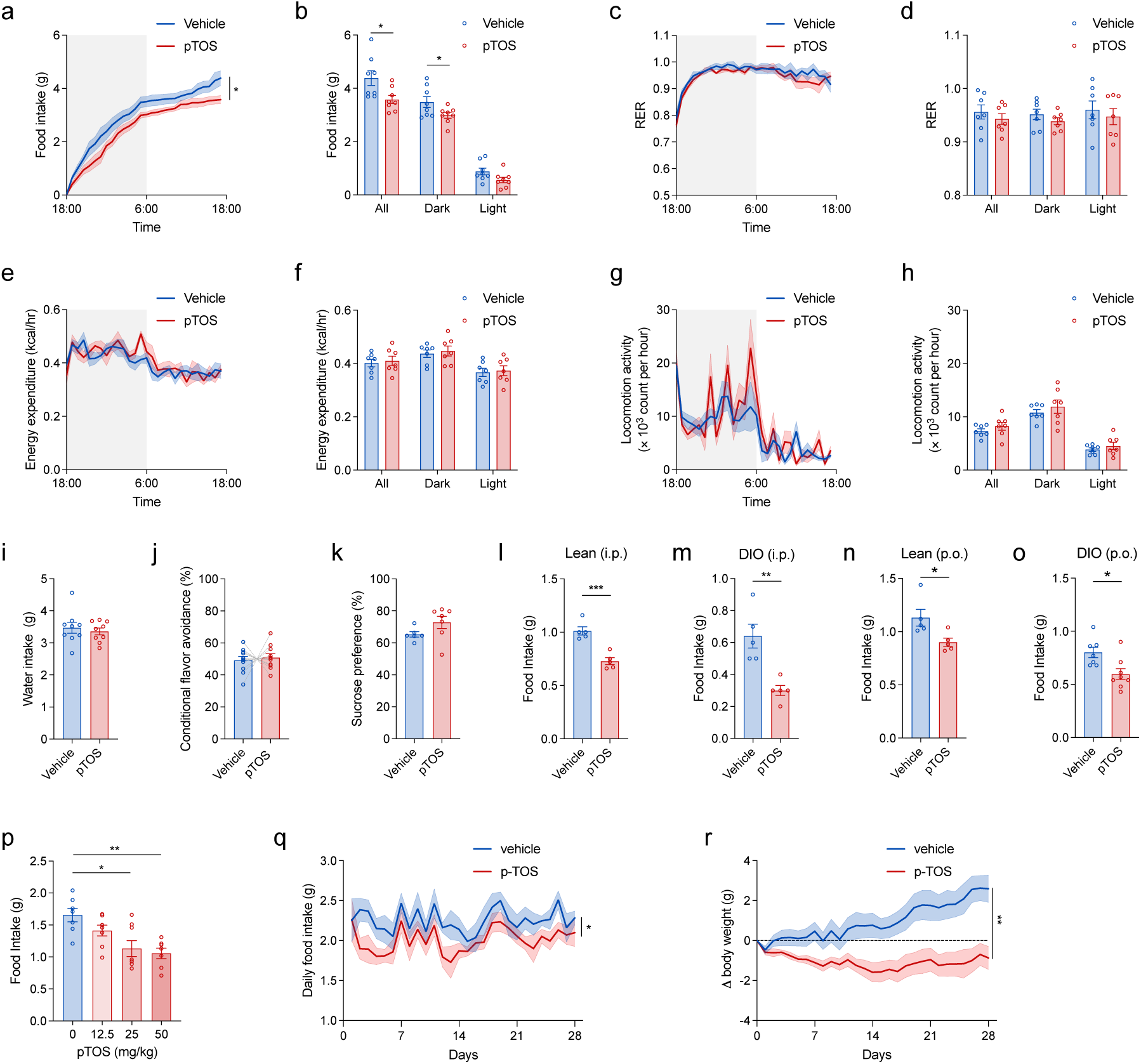
Effects of acute pTOS administration to mice. (a-h) Metabolic parameters measured with 24-hour fasted male mice (16-week-old) treated with pTOS (50 mg/kg, i.p.) or vehicle control. (a, b) Cumulative food intake (a) and food intake during the light and dark cycles (b) (n = 8/group). (c, d) Respiratory exchange ratio (RER) over time (c) and during the dark and light cycles (d) (n = 7/group). (e, f) Energy expenditure (adjusted for body mass) over time (e) and during the dark and light cycles (f) (n = 7/group). (g, h) Locomotor activity over time (g) and during the dark and light cycles (h) (n = 7/group). (i) Water intake (24 h) in male mice (16-week-old) treated with pTOS (50 mg/kg, i.p.) or vehicle control (n = 9/group). (j) Conditioned flavor avoidance ratio in paired solution for male mice (8-week-old) treated with pTOS (50 mg/kg, i.p.) or vehicle control. (n = 10/group). (k) Sucrose preference ratio in male mice (14-week-old) treated with pTOS (50 mg/kg, i.p.) (n = 7) or vehicle control (n = 6). (l, m) Three-hour food intake in lean (l, fed ad libitum, 12-14 weeks old) and diet-induced obese (DIO) male mice (m, fed ad libitum, 14-15 weeks old, 8-9 weeks on HFD, body weights were: vehicle 37.0 ± 2.5 g, pTOS 37.4 ± 0.7 g) treated with pTOS (50 mg/kg, i.p.) or vehicle control (n = 5/group). (n, o) Three-hour food intake in lean (n, n = 5, 11-13 weeks old) and DIO (O, n = 7 for vehicle, n = 8 for pTOS, 13-15 weeks old, 7-9 week on HFD, body weights were: vehicle 34.1 ± 1.2 g, pTOS 34.1 ± 1.0 g) male mice (fed ad libitum) treated with pTOS (50 mg/kg, p.o.) or vehicle control. (p) Three-hour food intake in lean (fed ad libitum, 14 weeks old) male mice treated with pTOS or vehicle (i.p.) at the doses indicated (n = 7 for 0 and 50 mg/kg, n = 8 for 12.5 and 25 mg/kg). (q, r) Daily food intake and body weight change of DIO mice (16 weeks old, 8 weeks on HFD, body weights were: vehicle 37.8 ± 3.3 g, pTOS 40.0 ± 3.0 g) with daily pTOS injections (50 mg/kg, i.p.) over a 28-day period. Data are shown as mean ± SEM. P values were calculated using two-way ANOVA (a, c, e, g, q, r), two-sided t tests (b, d, f, h, i, j, k, l, m, n, o), one-way ANOVA followed by Šídák’s multiple comparisons test (p), * p < 0.05. **p < 0.01, *** p < 0.001. Exact P values are as follows: (a) P = 0.041; (b) All, P = 0.0236, Dark, P = 0.0499, Light, P = 0.0741; (l) P = 0.0005; (m) P = 0.0031; (n) P = 0.0303; (o) P = 0.0150; (p) 0 vs 25, P = 0.0034, 0 vs 50, P = 0.0013; (q) P = 0.0494; (r) P = 0.0062.

We performed additional experiments to characterize the anorexigenic effect of pTOS in mice. In a 3-hr feeding experiment, pTOS (50 mg/kg, i.p.) reduced food intake in both lean (vehicle: 1.01 ± 0.04 g, pTOS: 0.73 ± 0.03 g, p = 5 × 10^-4^) and diet-induced obese (DIO; vehicle: 0.64 ± 0.07 g, pTOS 0.30 ± 0.03 g, p = 0.003) male mice housed in home cages (Figure 3l,m). Oral gavage administration of pTOS (50 mg/kg, p.o.) also increased plasma pTOS levels (Extended Data figure 5g) and reduced 3-hr food intake in both lean (vehicle: 1.13 ± 0.08 g, pTOS 0.90 ± 0.04 g, p = 0.030) and DIO (vehicle: 0.80 ± 0.05 g, pTOS: 0.60 ± 0.05 g, p = 0.015) male mice (Figure 3n,o). The higher exposure levels of pTOS following oral gavage versus IP administration may be due to its partitioning to fat/mesenteric tissues, which could slow its systemic release. The effect of pTOS on food intake was dose-dependent (Extended Data figure 5h and Figure 3p). Chronic administration of pTOS (50 mg/kg, i.p.) to DIO mice durably reduced daily food intake (vehicle: 2.25 ± 0.09 g, pTOS: 2.00 ± 0.07 g, p = 0.049) (Figure 3q) and produced a −9% vehicle-adjusted reduction in body weight (vehicle: 2.59 ± 0.68 g, pTOS: −0.87 ± 0.57 g, p = 0.002) (Figure 3r). We conclude that pTOS non-aversively reduces food intake and obesity without affecting water intake, energy expenditure, or movement.

Because pTOS is a sulfated tyramine analog, and because tyramine has been previously shown to suppress feeding^27,28^, we next sought to determine whether the anorexigenic activity of pTOS might be mediated by a tyraminergic activity of this metabolite. Using a cAMP-based cellular assay, we were unable to detect pTOS-dependent activation of the trace amine-associated receptor TAAR1^29^ (Extended Data figure 5i). In addition, administration of pTOS (50 mg/kg, i.p) to mice did not increase blood pressure (Extended Data figure 5j), whereas tyramine itself (50 mg/kg, i.p.) was, as expected, vasoactive in this assay (Extended Data figure 5k)^30^. We conclude that pTOS does not exhibit tyraminergic activity.

pTOS may also impact feeding by regulating the levels of other hormones previously implicated in food intake control. However, we did not detect any changes in plasma ghrelin (Extended Data figure 5l), leptin (Extended Data figure 5m), adiponectin (Extended Data figure 5n), insulin (Extended Data figure 5o), GLP-1 (Extended Data figure 5p), or GDF15 (Extended Data figure 5q) levels at either 1 h or 3 h after pTOS administration. pTOS did not affect systemic nutrient absorption following oral glucose, lipid, or protein administration (Extended Data figure 6a-f). pTOS also did not affect gastric emptying, whereas Exenatide, a GLP-1R agonist and positive control, showed the expected delayed emptying effect (Extended Data figure 6g). We conclude that the anorexigenic activity of pTOS is not dependent on changes in other known feeding-regulating hormones or on changes to gastric emptying or nutrient handling.

### VMH neurons mediate the anorexigenic activity of pTOS

Feeding behaviors are controlled centrally. To determine if pTOS might act directly in the brain, we measured metabolite levels in cerebrospinal fluid (CSF) and whole brain lysates following administration of either pTOS or tyramine control (50 mg/kg, i.p.) to mice. Unlike tyramine administration, which led to low peak plasma levels (Extended Data figure 6h) and low accumulation of tyramine in either CSF (Extended Data figure 6i) or brain lysates (Extended Data figure 6j), pTOS was robustly detected after pTOS administration both in CSF (at 12.6 μM, Extended Data figure 6k) and in total brain lysates (at 1.2 μM, Extended Data figure 6l) 30 min post-administration. We conclude that pTOS exhibits enhanced plasma stability and penetrates the central nervous system (CNS) to levels that may impact feeding pathways.

To understand the neural populations that might be modulated by pTOS, we performed activity mapping using the targeted recombination in active populations (TRAP) approach. We generated *TRAP2/Rosa26-LSL-tdTomato* mice^31^ (see Methods), which enable cFos-dependent recombination and genetic labeling of pTOS-activated neural populations with a tdTomato fluorescent reporter. *TRAP2/Rosa26-LSL-tdTomato* mice were treated with vehicle or pTOS (50 mg/kg i.p.) followed by 4-OHT (4-hydroxytamoxifen, 50 mg/kg, i.p.) to induce recombination. Two weeks later, we harvested the brains and examined multiple hypothalamic and brainstem regions for tdTomato^+^ signals (Extended Data figure 7). pTOS treatment increased the number of tdTomato^+^ cells in both the ventromedial hypothalamus (VMH) (Figure 4a,b) and the paraventricular hypothalamus (PVH) (Extended Data figure 7b). By contrast, the number of tdTomato^+^ cells was not statistically different between groups in the other brain regions examined (Extended Data figure 7c-m).

**Figure 4.**
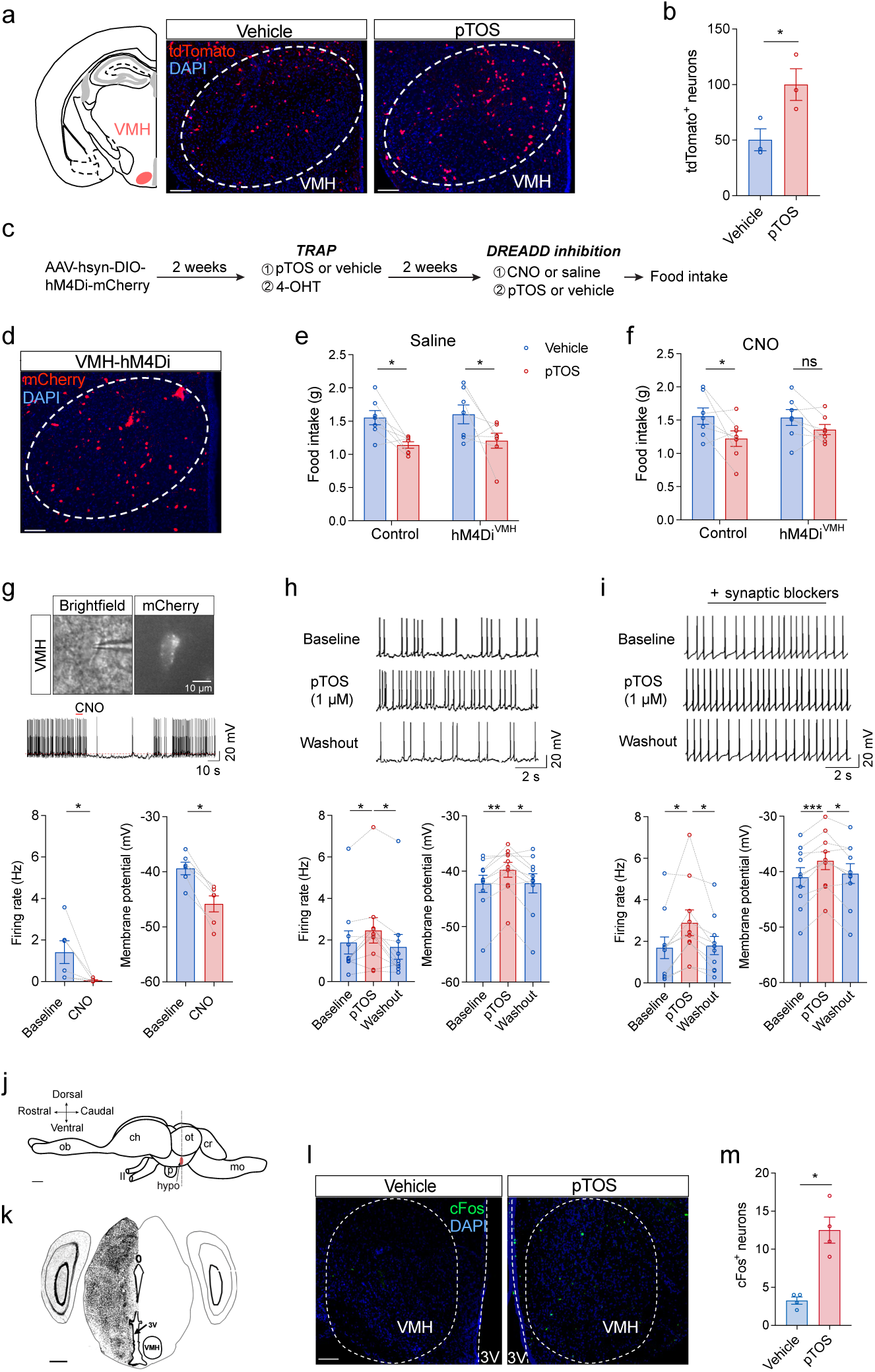
Role of VMH neurons in the anorexigenic effects of pTOS. (a) Anatomical location (left), and representative immunofluorescence section (right) of pTOS-activated neurons in the VMH in male *TRAP2/Rosa26-LSL-tdTomato* mice (10-week-old) following treatment with vehicle or pTOS (50 mg/kg, i.p., n = 3/group). VMH, ventromedial hypothalamus. (b) Quantification of pTOS-activated neurons in (a). (c) Schematic illustration of using hM4Di-DREADD to suppress pTOS-activated VMH neurons in TRAP2 mice. Cre-dependent AAVs carrying the hM4Di-mCherry DREADD were targeted to the VMH of TRAP2 mice for chemogenetic silencing. DREADD, designer receptors exclusively activated by designer drugs; 4-OHT, 4-hydroxytamoxifen; CNO, clozapine-*N*-oxide. (d) Representative image of hM4Di-mCherry expression in the VMH of male hM4Di^PVH^ mice, experiment repeated 7 times with similar results obtained. (e) Food intake (4-hour) in control or hM4Di^VMH^ mice pretreated with saline or CNO, followed with pTOS (50 mg/kg, i.p.) or vehicle administration (n = 7/group, 13-14 weeks old). (f) Representative image for electrophysiological recordings (top), representative action potential traces (middle), and quantitation of firing rate and resting membrane potential (bottom) of hM4Di^VMH^ neurons in response to CNO (10 µM, n = 6 neurons). (g, h) Representative action potential traces (top) and quantitation of firing rate and resting membrane potential (bottom) of hM4Di^VMH^ neurons following treatment with pTOS (1 µM) in the absence (g) or presence (h) of synaptic blockers (30 μM CNQX, 30 μM D-AP5, and 50 μM bicuculline) (n = 10 neurons/group) CNQX, cyanquixaline; D-AP5, 5-phosphono-D-norvaline. (j) Schematic drawing of the sagittal view of a ball python brain. Scale bar, 1 mm. Ch, cerebral hemisphere; cr, cerebellum; hypo, hypothalamus; mo, medulla oblongata; ob, olfactory bulb; ot, optic tectum; p, pituitary; II, second cranial nerve. (k) Coronal view of the brain section showing the VMH region. Scale bar, 500 µm. (l) Representative cFos staining in the VMH of fasted ball pythons treated with pTOS (50 mg/kg, p.o.) or vehicle control. Scale bar, 100 µm.(m) Quantitation of cFos^+^ neurons in the VMH of ball pythons treated with pTOS (50 mg/kg, p.o.) or vehicle control (n = 4/group). Data are shown as mean ± SEM. P values were calculated using two-sided t tests (b, m), one-sided paired t tests (e, f), two-sided paired Wilcoxon texts (g), one-way ANOVA followed by Turkey’s multiple comparisons tests (h, i),. * p < 0.05. **p < 0.01, *** p < 0.001. Exact P values are as follows: (b) P = 0.0464; (e) Control, P = 0.0070, hM4Di^VMH^, P = 0.0483; (f) Control, P = 0.0358, hM4Di^VMH^, P = 0.0929; (g) Firing rate, P = 0.0312, membrane potential, P = 0.0312; (h) Firing rate, Baseline vs pTOS, P = 0.0369, pTOS vs Washout, P = 0.0107, Membrane potential, Baseline vs pTOS, P = 0.0052, pTOS vs Washout, P = 0.0220; (i) Firing rate, Baseline vs pTOS, P = 0.0124, pTOS vs Washout, P = 0.0264, Membrane potential, Baseline vs pTOS, P = 0.0002, pTOS vs Washout, P = 0.0132; (m) P = 0.0020.

Next, we sought to determine whether the activation of VMH or PVH neurons was functionally relevant for the anorexigenic activity of pTOS. We therefore used viral approaches to enable chemical silencing of pTOS-activated neurons in these brain regions with the synthetic ligand clozapine-*N*-oxide (CNO) (Figure 4c). To first target pTOS-activated neurons in the VMH, we stereotaxically injected a Cre-dependent AAV carrying the inhibitory DREADD hM4Di-mCherry^32,33^ bilaterally into the VMH of *TRAP2* mice. Two weeks later, DREADD expression was induced in pTOS-activated VMH neurons (hM4Di^VMH^) by sequential administration of pTOS (50 mg/kg, i.p.) and 4-OHT (50 mg/kg, i.p.) (Figure 4d) As controls, similar procedures were performed in WT littermates. In an acute feeding assay, pTOS, as expected, suppressed food intake in three groups: control mice pretreated with saline (mean ± SEM, vehicle: 1.55 ± 0.11 g, pTOS: 1.14 ± 0.05 g, p = 0.007), control mice pretreated with CNO (vehicle: 1.56 ± 0.12 g, pTOS: 1.22 ± 0.12 g, p = 0.036), and hM4Di^VMH^ mice pretreated with saline (vehicle: 1.60 ± 0.14 g, pTOS: 1.20 ± 0.11 g, p = 0.048, Figure 4e). By contrast, the anorexigenic effect of pTOS administration was blunted in CNO-pretreated hM4Di^VMH^ mice (vehicle: 1.54 ± 0.12 g, pTOS: 1.36 ± 0.08 g, p = 0.093, Figure 4f). We conclude that pTOS-activated VMH neurons are required for the anorexigenic effects of pTOS.

To determine whether pTOS activates hM4Di^VMH^ neurons via a direct or indirect mechanism, we used post hoc slice electrophysiology to examine the activity of pTOS on these neurons in vitro. As a positive control, we first validated that in vitro treatment of hM4Di^VMH^ neurons with CNO reduced activity as expected (Figure 4g). Next, we observed that pTOS treatment (1 μM) of hM4Di^VMH^ neurons in vitro increased neuronal firing (vehicle: 1.88 ± 0.55 Hz; pTOS: 2.46 ± 0.61 Hz), an effect that was reduced following pTOS removal during the washout period (1.67 ± 0.60 Hz, Figure 4h). Furthermore, pTOS-dependent activation of hM4Di^VMH^ neurons persisted in the presence of a cocktail of synaptic blockers (Figure 4i). We conclude that pTOS directly activates hM4Di^VMH^ neurons in vitro.

We next turned to examine the role of pTOS-activated PVH neurons. We performed similar experiments to those described above, but via stereotaxic injection of hM4Di-mCherry virus bilaterally into the PVH of *TRAP2* or WT control mice, rather than the VMH (Extended Data figure 8a). In this experiment, pTOS administration equally suppressed food intake in all four groups of mice (Extended Data figure 8b,c). We confirmed the efficiency of hM4Di-mediated inhibition of hM4Di^PVH^ neurons using post hoc slice electrophysiology (Extended Data figure 8d-f). Thus, the targeted subset of PVH neurons, although activated by pTOS, are not required to mediate the effect of pTOS on feeding behaviors.

### Effect of pTOS on VMH neurons in pythons

Lastly, we sought to examine the effects of pTOS administration in pythons. Pythons eat in a single bout and consequently the rate of feeding is difficult to directly measure in these animals. As a surrogate molecular marker of pTOS activity, we examined brain cFos levels by immunofluorescence 90 min following administration of pTOS (50 mg/kg, p.o.) or vehicle to ball pythons. The brain of ball python is elongated along the sagittal plane, adapting to its narrow skull and predatory lifestyle (Figure 4j). The python VMH was easily identifiable as a distinct oval cluster of neurons located in the hypothalamus, near the third ventricle (Figure 4k)^34^. pTOS administration increased cFos staining in the VMH of ball pythons (Figure 4l,m). We conclude that pTOS activation of VMH neurons is a conserved activity across species.

In addition to the VMH, we also examined cFos staining following pTOS treatment in other brain regions as well. These other regions were identified using standard anatomical markers^34^. We observed that many other brain regions showed similar cFos immunoreactivity between vehicle and pTOS groups (Extended Data figure 9). We also observed that pTOS significantly inhibited neurons in the dorsal cortex (DC), piriform cortex (PC) and spinal trigeminal nucleus (SN) (Extended Data figure 9b,c,l). Therefore, pTOS-dependent activation of python VMH neurons is specific to neurons in that region, rather than reflecting a global increase in cFos activity following administration of this metabolite.

## Discussion

Pythons exhibit extreme feeding and fasting patterns and provide a unique model system for uncovering molecular regulators of the postprandial response. Using pythons as a discovery tool, here we provide multiple lines of evidence that pTOS is a postprandial anorexigenic metabolite that links nutrient intake to feeding control: 1) circulating pTOS levels are induced by >1,000-fold after feeding in pythons; 2) python pTOS can be produced from microbiome-dependent metabolism of dietary tyrosine; 3) administration of pTOS to pythons activates a neural population in the VMH, a key region that controls feeding and energy homeostasis; 4) pTOS-dependent activation of VMH neurons in mice drives suppression of food intake, demonstrating conserved activity across species; and 5) pTOS is detectable in the circulation in humans and its levels are also increased after a meal. The discovery of the postprandial induction and central activity of pTOS underscores the power of extreme model organisms to uncover molecules that might otherwise be missed in more classical organisms, such as mammals, which exhibit more limited fluctuations of energy states and postprandial physiology.

Many other gut-derived peptide and protein hormones have been previously studied in the context of mammalian postprandial metabolism, including CCK (cholecystokinin), GLP-1 (glucagon-like-peptide-1), GIP (gastric inhibitory peptide), PYY (peptide YY), and gastrin, and regulate diverse aspects of digestion, nutrient absorption, glucose regulation, and feeding^34–36^. A growing body of recent work also suggests that gut-derived metabolites such as bile acids and short-chain fatty acids can play similarly important roles^35–37^. Our data nominate pTOS as a gut-derived metabolite with a critical role in the python postprandial response. In the future, it will be important to understand the time course of pTOS induction relative to the other known postprandial peptide hormones in pythons. It will also be important in the future to determine if pTOS might also interact with any of these other peptide hormone pathways, including GLP-1. Lastly, we show that pTOS activates a neural population in the VMH, a region that is integral to many neuroendocrine functions including feeding and energy balance^38^. Projecting forward, the precise molecular identity of pTOS-activated VMH neurons, as well as the identity of specific downstream receptors within this cellular population that can be directly liganded by pTOS, still need to be identified. In addition, it is also possible that pTOS has additional bioactivity beyond feeding regulation alone.

Because pTOS is derived from dietary tyrosine, this metabolite can be considered a circulating sensor of tyrosine ingestion. It remains possible that other dietary components, metabolic stimuli, or endogenous secretions can also increase circulating pTOS levels. Currently the mechanisms responsible for the differing basal circulating levels of pTOS between Burmese and ball pythons, as well as humans, remains unknown, but could reflect differences in production, tissue distribution, catabolism, or clearance.

Chemically, pTOS is structurally related to tyramine, a monoamine metabolite and trace amine. Despite this structural similarity, sulfation here functions as a chemical switch that fundamentally alters the pharmacological and physiological properties of tyramine to both inactivate tyraminergic activity and to confer neo-activities to pTOS, such as enhanced plasma stability and improved brain penetration. This chemical switch mechanism for tyramine and pTOS parallels that of acetylation/methylation for serotonin and melatonin and hydroxylation for dopamine and norepinephrine. All these chemical modifications and metabolite pairs underscore how small chemical changes can lead to profound effects on downstream function. The possibility that sulfation modifications may function as a general molecular switch for other metabolites is an interesting possibility that remains largely unexplored.

While pythons consistently exhibit the most dramatic induction of pTOS after a meal, we also observe postprandial induction of pTOS in humans. On average, pTOS levels were increased by ∼2-5 fold following meals in most human cohorts examined. Nevertheless, we also found individuals with outlier pTOS responses: in the Mohold cohort study, one individual increased pTOS levels by >30-fold; in our own analysis of the Voldstedlund cohort study, another individual increased pTOS levels by >25-fold and reached python-level concentrations (∼2 µM at 0.5 h post-meal). Therefore postprandial regulation of pTOS is conserved in pythons and humans. In the future, it would be interesting to determine whether postprandial pTOS levels can be further augmented with larger, protein-enriched meals that more closely resemble the meals provided to pythons. It also remains possible that the observed difference in induction levels reflect other physiological changes following feeding, given that humans do not undergo extensive gastrointestinal remodeling. In addition, the absence of pTOS in mice was surprising, considering its conservation in other species. We speculate that mice might instead produce a chemically related, pTOS-like metabolite, analogous to how mice lack cortisol but use corticosterone as the functional equivalent^39^.

Lastly, this work builds on the long tradition of harnessing reptiles for natural products and drug discovery. Snake venoms, for example, have yielded a large number of bioactive molecules with clinical significance, including peptide modulators of blood pressure (e.g., ACE inhibitors and angiotensin receptor antagonists) and potent anti-thrombotic proteins (e.g., flavoridin and echistatin)^40–42^. Similarly, the Gila monster peptide Exendin-4 paved the way for the development of exenatide (Byetta), a first generation GLP-1 receptor agonist^43^. Although these previous discoveries primarily focused on snake venom, our data show that snake blood can also be a rich source of new molecular entities. Dynamic physiologic regulation, such as after a meal, can further help to identify high priority candidates for downstream functional testing.

## Materials and Methods

### Python husbandry

Animal protocols and procedures involving pythons were approved by the Institutional Care and Use Committee (IACUC) of the University of Colorado Boulder. Captive-bred Burmese pythons (*Python molurus bivittatus*) and Ball pythons (*Python regius*) were purchased from Bob Clark Reptiles, Oklahoma City, OK. Pythons were single-housed in the University of Colorado Boulder vivarium in 12-hour light-dark cycles at 30°C with 50% relative humidity. The pythons were then subjected to fasting and feeding schedules as in Figure 1a, consisting of a 28-day fasting period followed by feeding with a meal equal to 25% of their body weight. Burmese pythons were approximately 2 years old and weighed 1.5-2.5kg at the start of the study. Ball pythons were approximately 1 year old at the start of the experiments and weighed 400-500g.

### Mouse husbandry

All mouse experiments were performed according to procedures approved by the IACUC of Stanford University and Baylor College of Medicine. Mice were maintained in 12-hour light-dark cycles at 22 °C with 50% relative humidity. Mice were fed with a standard rodent chow diet (18% protein and 6% fat; Envigo Teklad 2018), or a high-fat diet (60% kcal fat; Research Diets, D12492), as specified.

### Acute mouse feeding studies

12-14 week-old *C57BL/6J* (Cat. 000664) and 14-15 week-old *C57BL/6J* DIO (Cat. 380050) male mice were obtained from The Jackson Laboratory. DIO mice were placed on high-fat diet for 8-9 weeks since 6 weeks old. Standard rodent chow (18% protein, 6% fat; Envigo Teklad 2018) or a high-fat diet (60% kcal from fat; Research Diets, D12492) were provided to lean or DIO mice, as specified. Mice were single-housed 3 hours prior to the onset of the dark cycle with ad libitum access to food and water. At the onset of the dark cycle, mice received a pTOS i.p. injection (50 mg/kg, or as indicated in figures) or oral gavage (p.o.). Food weight was recorded at baseline (0 hr) and after 3 hr. The body weights of DIO mice were: i.p. vehicle 37.0 ± 2.5 g, pTOS 37.4 ± 0.7 g; p.o.: vehicle 34.1 ± 1.2 g, pTOS 34.1 ± 1.0 g.

### Chronic pTOS treatment in DIO mice

For the chronic treatment, wild-type *C57BL/6J* male mice (8 weeks old) were fed with a high-fat diet (60% kcal from fat; Research Diets, D12492) for 8 weeks. Initial body weights of the mice were: vehicle 37.8 ± 3.3 g, pTOS 40.0 ± 3.0 g. The mice were then single housed and administrated pTOS (50 mg/kg, i.p.) or vehicle daily three hours before the onset of the dark cycle for 28 executive days. The mixture of 18:1:1 (by volume) of saline:Kolliphor EL (C5135, Sigma):DMSO was used as vehicle. Daily food weight and body weight were recorded.

### Python feeding studies

Pythons were randomly assigned to different endpoint groups. They were then fasted for 28 days, after which they received a rat meal equivalent to 20-25% of their body weight. At each pre-assigned endpoint, pythons were euthanized via rapid decapitation under deep isoflurane-induced anesthesia. Sufficient anesthetic depth was confirmed by lack of response to physical stimuli. Blood was collected in BD Vacutainer Lithium Heparin tubes (Fisher Scientific), immediately mixed by inversion, placed on ice for 10 minutes, and then plasma was separated by centrifugation at 3,000 g for 15 minutes to pellet red blood cells. The supernatant was collected, flash-frozen in liquid nitrogen, and stored at −80°C until further analysis. Dissected tissues were rinsed in ice-cold PBS and then flash frozen in liquid nitrogen and stored at −80°C until further analysis.

### Python oral gavage studies

Ball pythons were randomly assigned to experimental or control groups and then fasted for 28 days. Tyrosine, water, or pTOS were then delivered by oral gavage using a 5 Fr. 16-inch rubber Sovereign Sterile Feeding Tube (MWI). Tyrosine was administered at 1 g/kg dissolved in water. pTOS was administrated at 50 mg/kg in an 18:1:1 mixture of water: DMSO: Kolliphor. Pythons that received pTOS or vehicle control were euthanized and dissected 90 minutes post-gavage. Pythons that received tyrosine were euthanized and dissected 24 hours post-gavage. Plasma and tissues were collected as described above.

### Human meal test study (Voldstedlund Study)

Arterial plasma samples were obtained from a meal test study that was approved by the Research Ethics Committee of Copenhagen and previously published^24^. Briefly, ten young male human subjects (age: 26.7 ± 1.3 years; BMI: 22.6 ± 0.6 kg/m^2^) first performed 60 min one-legged dynamic knee extensor exercise 90 min after a small breakfast (18kJ per kg) in the morning. Four hours later (time “0 min”), the subjects received a mixed solid meal (30 kJ per kg body weight), followed by two mixed liquid meals (20 kJ per kg body weight; Nutridrink, Nutrica, Denmark) 30 min and 60 min after the solid meal. The mixed meals were composed of (energy intake) 50% carbohydrates, 35% fat, and 15% protein.

### Tyramine production from fecal samples

Fecal samples from ball pythons (21 mg) were dispensed in 500 μL Standard Amino Acid Complete (SAAC) media, previously described^44^. The mixture was centrifuged at 100 g for 5 min to pellet undigested materials. 100 μL of the supernatant was mixed with 300 μL SAAC media and incubated in an anaerobic chamber (Coy Laboratories) at 37 °C for 2 days, in an atmosphere of 5% hydrogen, 10% carbon dioxide, and 85% nitrogen. Control SAAC medium contained 1 mM tyrosine and 0.1 mg/L vitamin B6. Testing medium contains 50 mM tyrosine and 5 mg/L vitamin B6. Cells were pelleted by centrifugation at 3,381 x g to separate bacteria from the conditioned media. 150 μL of 2:1:1 acetonitrile: methanol: water mixture was added to extract metabolites from the bacteria and 50 μL of media were mixed with 150 μL 2:1 acetonitrile: methanol to extract metabolites from media.

### Tyramine production from python microbiome

Intestinal content was harvested from the small and large intestines of a fed (3DPF) and a fasted (107DPF) python and stored at −80°C. The intestinal content was then washed in reduced PBS supplemented with 0.05% cysteine, and centrifuged at 100 g for 5 min to pellet undigested materials. The supernatant was then centrifuged at 5,000 g for 5 min to pellet bacteria. Bacteria were resuspended in 10 mL of reduced Gifu media. A final concentration of 10 g/L tyrosine and 500 mg/L vitamin B6 was used in the experimental group. The bacteria were cultured for 48 hours at 30°C in an anaerobic chamber. At the end of the incubation period, the cells were pelleted by centrifugation at 5,000 g for 5 minutes. The media was collected for further LC-MS analysis.

### Python antibiotic treatment

A cocktail of antibiotics (ABX) containing 200 mg/kg metronidazole, 200 mg/kg ampicillin, 100 mg/kg neomycin, and 100 mg/kg erythromycin was delivered to juvenile Burmese pythons (weighing 125-200 g at time of study) via oral gavage over a seven-day period. The ABX cocktail was administrated to pythons via oral gavage on day 1, 2, 3, 5, and 7. Control pythons received oral gavage of vehicle (water) at an equivalent volume. Concurrently, ABX-treated pythons also received metronidazole, ampicillin, neomycin, and erythromycin (each at 0.1g/L) in the drinking water. Seven days after the first dose of antibiotics, pythons were either euthanized or fed a mouse meal equal to 25% of their body weight. Antibiotic-treated pythons received a germ-free (GF) mouse meal, whereas vehicle-treated pythons received a specific pathogen-free (SPF) mouse meal. ABX-treated pythons continued to receive the antimicrobial cocktail via drinking water after their meal. Pythons were euthanized with or without ABX treatment at 3- and 28- days post feeding (DPF). Plasma was collected as above.

### pTOS production with python liver slices

Fed (3DPF) and fasted (31DPF) ball pythons were euthanized via rapid decapitation while under anesthesia and the liver was immediately collected and washed in sterile PBS. Freshly isolated liver was transferred to a sterile fume hood and 0.5 g of total liver mass was diced into fine segments (∼2×2 mm) and then placed into one well of a 6-well plate containing 2mL William’s E media supplemented with 1% non-essential amino acids (NEAA), 1% GlutaMAX, 2% fasted python plasma, 100 nM dexamethasone, 100 nM insulin, and 0.375% fatty-acid free BSA. Tyramine (1 mM final concentration) or vehicle control was added to the media and incubated at 30°C, 5% CO_2_ for 16 hours. The media was collected and clarified by centrifugation at 21,130 x g for 10 minutes. The supernatant was flash frozen in liquid nitrogen and stored at −80°C until further analysis.

### Python sulfotransferase expression in HEK293T cells and enzymatic assay

Aryl sulfotransferases (SULTs) with higher abundance in postprandial python livers were codon-optimized for expression in HEK293T cells and synthesized by IDT. The sequences were then cloned into a pCMV5-mCherry vector with a 3x FLAG tag on the C-terminal. For simplicity, we named the python SULTs with the following annotations based on the sequence similarities: SULT1A1 (A0A9F5IVD0), SULT1C4 (A0A9F2QWK4), SULT1D1 (A0A9F2REM3), and SULT6B1 (A0A9F2R4U1). HEK293T cells were transiently transfected with Polyfect. 24 hours later, the cells were washed with ice-cold PBS and scraped into an Eppendorf tube. The cells were pelleted by centrifugation at 845 g for 5 minutes at 4°C, and lysed in 50 mM potassium phosphate buffer (pH 7.4) by sonication. The protein concentration of the soluble fraction was determined by BCA assay. The tyramine sulfotransferase activity was determined with cell lysates containing 100 μg total protein, supplemented with 100 μM tyramine, 100 μM PAPS (R&D Sytems, ES019), and 1 mM pargyline (Cayman, 10007852), a MAO inhibitor. The assay was initiated by incubation at 30°C for 30 min and stopped by adding ice-cold acetonitrile:methanol. SULT expression was validated by Western blot. SULT1D1 was not expressed in HEK293T cells, due to its incomplete sequence in Uniprot. The activity was also indistinguishable from non-transfected controls. Heat-inactivated cell lysates exhibited non-detectable enzymatic activity.

### Tyramine and pTOS extraction from cultured media for LC-MS

To extract metabolites from the media of cultured hepatocytes or bacteria, 200 µL of 1-butanol was added to 500 µL media. The mixture was vortexed for 1 minute, and then centrifuged at 4 °C for 10 minutes at 21,130 x g. The top layer was carefully transferred to mass spec vials for LC-MS analysis.

### Mouse blood and plasma sample preparation for LC-MS

Blood from mice was collected by submandibular bleeding into lithium heparin tubes (BD, 365985) and immediately kept on ice. The blood was then centrifuged at 4 °C for 5 min at 2,348 x g. Plasma was transferred into new Eppendorf tubes and stored at −80 °C if not immediately used. Metabolites were then extracted by adding 150 µL of a 2:1 mixture of acetonitrile: methanol to 50 µL of serum or plasma.

### LC-MS analysis

Untargeted metabolomics measurements were performed using an Agilent 6545 Quadrupole time-of-flight (QTOF) LC/MS instrument. MS analysis was performed using electrospray ionization (ESI) in both positive and negative modes. The dual ESI source parameters were set as follows: the gas temperature was set at 250 °C with a drying gas flow of 12 L/min and the nebulizer pressure at 20 psi; the capillary voltage was set to 3,500 V; and the fragmentor voltage set to 100 V. Separation of metabolites was conducted using a Luna 5 μm NH_2_ 100 Å LC column (Phenomenex, 00B-4378-E0) with normal phase chromatography. Mobile phases in positive mode were: buffer A, water with 0.1% formic acid; buffer B, acetonitrile with 0.1% formic acid. Mobile phases in negative mode were: buffer A, 95:5 water:acetonitrile with 0.1% ammonium hydroxide and 10 mM ammonium acetate; buffer B, acetonitrile. The LC gradient started at 100% buffer B with a flow rate of 0.7 mL/min from 0 to 2 min. The gradient was then linearly increased to 50% buffer A and 50% buffer B at a flow rate of 0.7 mL/min from 2 to 20 min. From 20 to 25 min, the gradient was maintained at 50% buffer A and 50% buffer B at a flow rate of 0.7 mL/min.

Quantification of the metabolite concentrations was performed by generating a standard curve with known concentrations of each metabolite. Metabolite standards were analyzed alongside the samples using the same method. A standard curve generated from the metabolite concentrations and extracted ion intensities was used to calculate the concentrations of each metabolite.

### Blood pressure measurements

Mice were anaesthetized with isofluorane and placed in the supine position with the fur removed in the chest area. Blood pressures were measured with a 1.4-F pressure sensor mounted Millar catheter (SPR-671, ADInstruments) that was inserted into the right carotid artery. Blood pressures were recorded with LabChart 7 Pro (ADInstruments) and annotated with the BP_annotate package in Matlab^45^.

### TSE PhenoMaster metabolic studies in mice

Sixteen-week-old *C57BL6/J* male mice were acclimated into the TSE PhenoMaster Metabolic Cage system. In the TSE PhenoMaster cages, mice were maintained on a chow diet for the first 3 days and then fasted for 24 hours followed by injection with vehicle or pTOS (50 mg/kg, i.p.) and refeeding at 5:30 pm before the start of the night cycle, while food intake, RER, energy expenditure, and locomotion were continuously monitored. Energy expenditure data were analyzed with each animal’s body weight as a covariate using the online CalR tool^46^. The mixture of 18:1:1 (by volume) of saline:Kolliphor EL (C5135, Sigma):DMSO was used as vehicle. pTOS was dissolved in the mixture, then aliquoted and stored at −80 °C before use.

### TRAP2 mice

We crossed *TRAP2* mice (Jackson Laboratory, #030323) with *Rosa26-LSL-tdTomato* mice (Jackson Laboratory, #007905) to generate *TRAP2/Rosa26-LSL-tdTomato* or *TRAP2* mice. Mice were housed in a temperature-controlled environment using a 12-hour light and 12-hour dark cycle. Mice were individually housed at least 1 week before the study. The mice were fed a standard chow diet (19.0% protein, 6.5% fat, 2.7% crude fiber, 12.3% neutral detergent fiber, by weight, Harlan-Teklad, Madison, Wisconsin, #2920). Water was provided ad libitum.

### TRAP induction

We dissolved 4-hydroxytamoxifen (4-OHT) (Sigma, Cat# H6278) at 20 mg/ml in ethanol by sonication at 37 °C for 15 min. The dissolved 4-OHT was then stored in aliquots at - 80 °C for up to several weeks or used immediately. Before use, 4-OHT was dissolved by shaking at 37 °C for 10 minutes, then sunflower seed oil and castor oil (4:1) was added for a final concentration of 10 mg/ml. After evaporating the ethanol in a vacuum (845 x g, 15 minutes), the final 4-OHT solution was injected intraperitoneally (i.p.) at 50 mg/kg. To TRAP pTOS-activated neurons, *TRAP2/Rosa26-LSL-tdTomato* male mice (10 weeks of age) were fasted from 12 pm to 5 pm, then received i.p. injection of pTOS (50 mg/kg), followed by 4-OHT injection (50 mg/kg, i.p.) 30 min after; two weeks later, the mice were perfused with saline followed by 10% formalin. As controls, another group of *TRAP2/Rosa26-LSL-tdTomato* male mice received i.p. injection of vehicle, followed by 4-OHT injection (50 mg/kg, i.p.) 30 min after; two weeks later, the mice were perfused. The mixture of 18:1:1 (by volume) of saline:Kolliphor EL (C5135, Sigma):DMSO was used as vehicle. Coronal brain sections were cut at 30 μm and collected into five consecutive series. Sections were cover-slipped and analyzed using a fluorescence microscope. The numbers of tdTomato-labeled (TRAPed) neurons were counted and quantified manually. Briefly, for each mouse brain structure analyzed, anatomically defined ROIs were selected based on the Allen Mouse Brain Atlas. The same ROI boundaries were consistently applied across all mice and sections. To ensure consistency, a single coronal brain section per region was analyzed for each mouse, selected at matched anterior-posterior coordinates based on anatomical landmarks. Three mice were included in each group.

### Chemogenetics approaches

Male *TRAP2* mice (8 weeks of age) were anesthetized (with 2% isoflurane) and placed in a stereotaxic instrument. Artificial eye ointment was applied to prevent corneal drying, and a heat pad was used to hold the body temperature at 37°C. To chemogenetically inhibit pTOS-activated PVH or VMH neurons, we injected AAV8-DIO-hM4Di-mCherry (Addgene, #44362, titer: 5×10^12^ GC/ml, 0.2 µl) into the PVH or VMH of TRAP2 mice (PVH: −0.82 mm; ML: 0.25 mm; DV: −4.75 mm; VMH: −1.7 mm; ML: 0.3 mm; DV: −5.6 mm), respectively. After allowing 2 weeks for virus expression, mice were singly housed before they were subjected to any studies. Mice were fasted overnight (from 5 pm to 9 am), then received saline or CNO (i.p., 3 mg/kg, #16882, Cayman) injections. After saline or CNO injection, pre-weighed regular chow (6.5% fat; #2020, Harlan Teklad) was put back into the cages. Food intake was monitored for 3 h. At the end of the experiments, all mice were perfused with saline followed by 10% formalin. Brain sections were collected and sectioned at 30 µm. The expression of mCherry was examined with histology, and only those with accurate targeting were included for data analyses.

### cFos mapping of pTOS-activated neurons in python brains

To process python brains, we adapted the approach as previously described for *in vivo* fixation of the lizard brain^48^. One hour after delivery of 50 mg/kg pTOS by oral gavage, Ball pythons were anesthetized via isoflurane inhalation for 30 minutes. Sufficient anesthetic depth was determined by lack of response to a physical stimulus. The thoracic cavity and rostral ∼5-cm were then opened with surgical scissors to expose the heart and carotid arteries. Fine incisions were made in the ventricle and right atrium, and a perfusion needle was inserted into the ventricular incision, through the right aorta, and into the left carotid artery. The needle was clamped in place with a hemostat and 50 mL of heparinized PBS (10 units/L) was perfused using a peristaltic pump to clear blood from the brain. Brain fixation was performed by perfusion with 50 mL of 4% paraformaldehyde (PFA). The brain was then extracted using a corneoscleral punch and submerged overnight in 4% PFA at 4°C. The brain was removed from PFA, carefully rinsed twice with PBS, and then added to a 30% sucrose PBS solution and stored at 4°C for two days to dehydrate the tissue. Coronal brain sections were cut at 30 µm and collected into five consecutive series. One series of the sections was blocked for 1 hour in 0.3% PBST with 5% normal donkey serum. To detect cFos expression, the Rabbit anti-cFos antibody (1:500, #226008, Synaptic System) was added and incubated at 4 °C overnight on shaker. The cFos antibody recognize the amino acid sequence “MFSGFNADYEASSSR”. Three out of the 15 amino acids in this sequence differ from those in the corresponding sequence in the python sequence. The following day, slices were rinsed with 0.1% PBST for 6 x 10 minutes and then incubated with donkey anti-rabbit AlexaFluor 488 (1:500, A21206, Invitrogen) at room temperature for 2 hours. Sections were cover-slipped and analyzed using a fluorescence microscope. The numbers of cFos-labelled neurons were counted and quantified manually. ROIs for each ball python brain structure were selected using the most reliable anatomical landmarks available from the limited existing references. For each ball python, 3-4 coronal brain sections per VMH were analyzed. Four pythons were included in each group. Locations of cFos^+^ neurons in the VMH and other brain regions were mapped onto the standard anatomical brain sections of *Python regius*^34^. Furthermore, brain sections from both the same species of red-sided garter snake^49,50^ and different species with similar brain structures, such as the tree lizard *Urosaurus ornatus*^51^, reveal consistent VMH locations. Within the Squamata order, the VMH is one of the brain regions existing the least variation, as determined by calculating the log-transformed coefficient of variation for each brain region to assess individual differences^52^.

### Statistics

The minimal sample size was pre-determined by the nature of the experiments. For biochemical measurements, at least 3-4 different mice or pythons per group were used. For behavioral measurements, 7-9 different mice per group were included. For histology studies, the same experiment was repeated in at least 3 different mice or pythons. For electrophysiological studies, at least 6 different neurons from 3 different mice were included. The data are presented as mean ± SEM or as individual data points. Statistical analyses were performed using GraphPad Prism to evaluate normal distribution and variations within and among groups. Methods of statistical analyses were chosen based on the design of each experiment and are indicated in figure legends.

## Data availability

All data generated or analyzed during this study are included in this article and its Supplementary Information files.

## Materials & Correspondence

Requests should be directed to

## Acknowledgements

We thank members of the Long lab for discussions. We thank Bernadette Gallardo for assistance in laboratory operations. We thank Dr. Jonathan Van Vranken for assistance with the multiplex proteomics at Harvard Medical School. We thank Haley Lavach in the BioFrontiers Institute at the University of Colorado for the expert help with python microbiome studies. This work was supported by the US National Institutes of Health (R01GM029090 to L.A.L., R01DK138518 to Y.X., R01DK105203 and R01DK124265 to J.Z.L., K99DK141966 to S.X., K99AR081618 to M.Z., F32HD112123 to M.W., F32HL170637 to T.G.M., F32DK138685 to X.F., T32GM142607 to M.P.M.), the Wu Tsai Human Performance Alliance (research grant to J.Z.L., postdoctoral fellowship to S.X.), the Stanford Diabetes Research Center (P30DK116074 to J.Z.L.), the Phil and Penny Knight Initiative for Brain Resilience at the Wu Tsai Neurosciences Institute (research grant to J.Z.L.), the Ono Pharma Foundation (research grant to J.Z.L.), the Leducq Foundation (21CVD02 to L.A.L.), the American Heart Association (24POST1200064 to S.X., 24POST1196199 to W.W.), Stanford University Medical Scientist Training Program (T32-GM007365 to S.D.T). The clinical trial of the Moholdt Study was supported by Novo Nordisk Foundation (NNF14OC0011493 to J.A.H.) and The Liaison Committee for Education, Research and Innovation in Central Norway (2016/29014 to T.M.).

## Author contributions

Conceptualization: S.X. and J.Z.L. Investigation: S.X., M.W., T.G.M., B.S. X.F., X.L., Y.Y., S.F., S.D.T., J.F.G., G.L.M., M.P.M., J.H.H., V.L.L., A.L.M., M.Z., W.Q., S.C.R., M.Z., J.S., W.W., T.M., C.T.V. Writing original draft: S.X. and J.Z.L. Writing, reviewing and editing: S.X. M.W., T.G.M., L.A.L., Y.X., and J.Z.L. Resources: J.A.H., E.A.R., X.C., K.J.S., D.B., L.A.L., Y.X., J.Z.L. Supervision and funding acquisition: L.A.L., Y.X., J.Z.L.

## Supplementary tables

Table 1. Plasma metabolomics of Fasted and 3DPF Burmese pythons.

Table 2. Liver proteomics of Fasted, 1DPF, and 3DPF ball pythons.

